# The *Coxiella burnetii* secreted protein kinase CstK influences vacuole development and interacts with the GTPase-activating protein TBC1D5

**DOI:** 10.1101/611707

**Authors:** Solene Brelle, Eric Martinez, Sylvaine Huc-Brandt, Julie Allombert, Franck Cantet, Laila Gannoun-Zaki, François Letourneur, Matteo Bonazzi, Virginie Molle

## Abstract

*Coxiella burnetii* is the etiological agent of the emerging zoonosis Q fever. Crucial to the pathogenesis of this intracellular pathogen is the secretion of bacterial effectors into host cells by a Type 4b Secretion System (T4SS), to subvert host cell membrane trafficking, leading to the biogenesis of a parasitophorous vacuole allowing intracellular replication. The characterization of prokaryotic Serine/Threonine Protein Kinases (STPKs) in bacterial pathogens is emerging as an important strategy to better understand host-pathogen interactions. In this study, we investigated CstK (for *Coxiella* Ser/Thr kinase), a bacterial protein kinase identified in *C. burnetii* by *in silico* analysis. Here, we demonstrated that this putative protein kinase undergoes autophosphorylation on Ser, Thr, and Tyr residues, and phosphorylates a classical eukaryotic protein kinase substrate *in vitro*. This dual Ser/Thr and Tyr kinase activity is similarly observed for eukaryotic dual specificity Tyr phosphorylation-regulated kinase class. CstK is translocated during infections and localizes at *Coxiella*-containing vacuoles (CCVs). Moreover, a *C. burnetii* mutant strain overexpressing CstK displays a severe CCVs development phenotype, suggesting a finely tuned regulation by the bacterial kinase during infection. Protein-protein interaction studies identified the Rab7-GTPase activating protein (GAP) TBC1D5 as a candidate CstK-specific host target, suggesting a role for this eukaryotic GAP in *Coxiella* infections. Indeed, CstK colocalizes with TBC1D5 in non-infected cells, and TBC1D5 is recruited at CCVs during infection. Accordingly, depletion of TBC1D5 from infected cells significantly affects CCVs development. Our results indicate that CstK has a critical role during infection as a bacterial effector protein that interacts with host proteins to facilitate vacuole biogenesis and intracellular replication.

---

Signal transduction is an essential and universal function that allows all cells, from prokaryotes to eukaryotes, to translate environmental signals to adaptive changes. By this mechanism, extracellular inputs propagate through complex signaling networks whose activity is often regulated by reversible protein phosphorylation. Signaling mediated by Serine/Threonine/Tyrosine protein phosphorylation has been extensively studied in eukaryotes, however, its relevance in prokaryotes has only begun to be appreciated. The recent discovery that bacteria also use Ser/Thr/Tyr kinase-based signaling pathways has opened new perspectives to study environmental adaptation, especially in the case of bacterial pathogens, with respect to host infection. Thus, advances in genetic strategies and genome sequencing have revealed the existence of “eukaryotic-like” serine/threonine protein kinases (STPKs) and phosphatases in a number of prokaryotic organisms, including pathogens such as, *Streptococcus* spp. (1–4), *Mycobacteria* (5–10), *Yersinia* spp. (11,12), *Listeria monocytogenes* (13,14), *Pseudomonas aeruginosa* (15), *Enterococcus faecalis* (16) or *Staphylococcus aureus* (17–20). Consequently, the study of STPKs in human bacterial pathogens is emerging as an important strategy to better understand host-pathogen interactions and develop new, targeted antimicrobial therapies. However, if on one hand it is clear that STPKs and phosphatases regulate important functions in bacterial pathogens, their signal transduction mechanism remains ill-defined and restricted to a limited number of microbes.

Importantly, STPKs expressed by pathogenic bacteria can either act as key regulators of important microbial processes, or be translocated by secretion systems to interact with host substrates, thereby subverting essential host functions including the immune response, cell shape and integrity (21). Phosphorylation of host substrates has been demonstrated for some bacterial STPKs, whereas others seem to require their kinase activity but their phosphorylated substrates remain to be identified (21). Therefore, biochemical mechanisms of these pathogen-directed targeted perturbations in the host cell-signaling network are being actively investigated and STPKs are proving to be molecular switches that play key roles in host-pathogen interactions (21).

Among emerging human pathogens, *Coxiella burnetii* is a highly infectious bacterium, responsible of the zoonosis Q fever, a debilitating flu-like disease leading to large outbreaks with a severe health and economic burden (22–24). The efficiency of infections by *C. burnetii* is likely associated with the remarkable capacity of this bacterium to adapt to environmental as well as intracellular stress. Indeed, outside the host, *C. burnetii* generates pseudo-spores that facilitate its airborne dissemination. During infection, *C. burnetii* has developed a unique adaptation to the host, being the only bacterium that thrives in an acidic compartment containing active lysosomal enzymes. Similarly to other intracellular bacterial pathogens, *C. burnetii* adaptation depends on the translocation of bacterial effectors by a Dot/Icm Type 4b Secretion System (T4SS). Some of these effectors modulate important signaling pathways of infected cells, including apoptosis and inflammasome activation (25–27), whereas others localize at *Coxiella*-containing vacuoles (CCVs) and manipulate host membrane traffic to facilitate the development of the intracellular replicative niche (28,29). *C. burnetii* genome analysis revealed a close homology to the facultative intracellular pathogen *Legionella pneumophila*, in particular at the level of Dot/Icm core genes (30). *In silico* analysis identified over 100 candidate effector proteins encoded in the *C. burnetii* genome, some of which have been validated for secretion using either *C. burnetii* or *L. pneumophila* as a surrogate model (31–33). In this study, we investigated the candidate effector CBU_0175, which encodes the only putative *Coxiella* Ser/Thr kinase (CstK). We demonstrated CstK translocation by *C. burnetii* during infection and we reported its localization at CCVs. *In vitro* kinase assays revealed that CstK undergoes autophosphorylation on Ser, Thr, and Tyr residues, and CstK displays kinase activity towards a test substrate of eukaryotic protein kinases. Furthermore, the identification of the Rab7 GTP-activating protein (GAP) TBC1D5 as a CstK interactor suggests that this protein might be phosphorylated during infections to facilitate CCVs biogenesis. Indeed, TBC1D5 is actively recruited at CCVs during *Coxiella* infections and TBC1D5-targeting siRNAs significantly affect CCVs development. Our data provide the first evidence that a *C. burnetii* secreted kinase might control host cell infection.

## RESULTS

### Coxiella burnetii genome encodes a single putative protein kinase

*In silico* analysis of the epidemic *C. burnetii* strain RSA493 NMI genome revealed only one gene encoding a putative Serine/Threonine Protein Kinase (STPK). To date, no STPKs have been characterized in this organism. This gene was named *cstK* for *C. burnetii* <u>s</u>erine <u>t</u>hreonine <u>K</u>inase and encodes a 246 amino acids protein with an estimated molecular mass of 31 kDa. ClustalW and BLAST analysis revealed the presence of most of the essential amino acids and sequence subdomains characterizing the Hanks family of eukaryotic-like protein kinases (34). CstK shares a common eukaryotic protein kinases superfamily fold with two lobes and a Gly-rich loop. These include the central core of the catalytic domain, and the invariant lysine residue in the consensus motif within subdomain II, which is usually involved in the phosphotransfer reaction and required for the autophosphorylating activity of eukaryotic STPKs (Fig. 1A) (34,35). The activation loop in the catalytic domain is particularly short in CstK, and the DFG motif is substituted by a GLG motif. Interestingly, the transmembrane domain usually presents in classical prokaryotic STPKs is lacking in CstK, thus it is a so-called cytoplasmic STPK.

**Figure 1.**
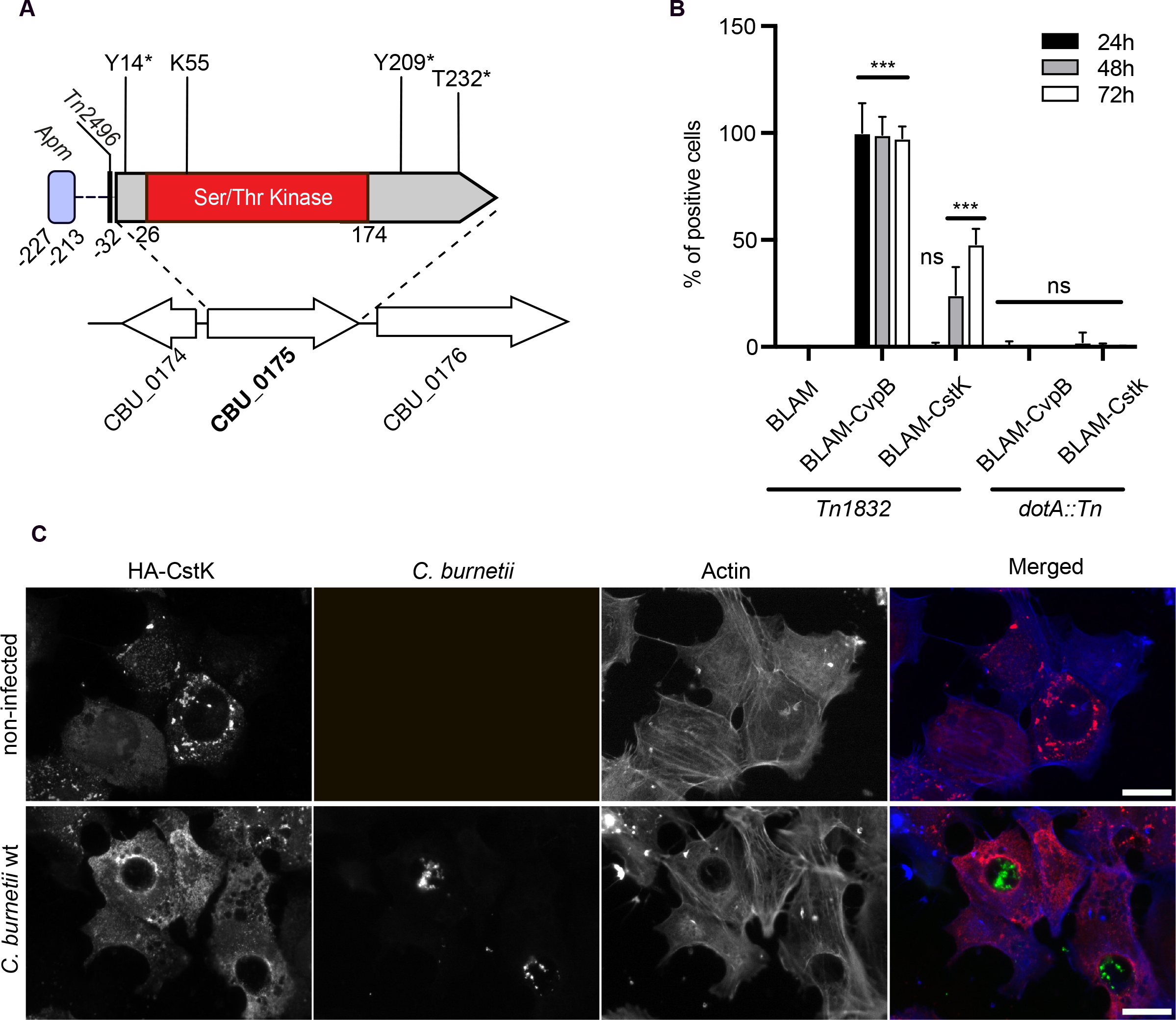
CstK is a Dot/Icm effector protein. **(A)** schematic representation of CstK. The kinase domain is shown in red, the predicted E-block for effector protein translocation in green. A conserved promoter-binding domain is predicted between aa −227 and −213 (*Apm*) and the transposon insertion site for *Tn2496* is indicated at 32 aa upstream of the *cstK* starting codon. The conserved lysine residue is indicated, and the phosphorylated sites indicated by an asterisk. **(B)** U2OS cells were infected for 24, 48, or 72h (black, grey and white bars, respectively) with *C. burnetii Tn1832* (wt) or the *dotA*::Tn mutant transformed with vectors expressing Beta-Lactamase alone (BLAM, negative control), BLAM-CvpB (positive control) or BLAM-CstK. Protein translocation was probed using the CCF-4 substrate. The average percentage of cells positive for cleaved CCF-4 as compared to the total number of cells was assessed. Values are mean ± SD from 3 independent experiments. ns = non-significant; *** = P<0.0001, one-way ANOVA, Dunnett’s multiple comparison test. **(C)** U2OS cells ectopically expressing HA-tagged CstK (red) were either left untreated (top panels) or challenged with *C. burnetii Tn1832* (wt) for 3 days. Actin (blue) was labelled with Alexa647-tagged phalloidin. Scale bars 10 μm.

### CstK is a Dot/Icm effector protein

Bioinformatics analysis using the prediction software S4TE2 (36) indicated that CstK harbors features corresponding to secreted effector proteins, including a promoter motif typically found in effector proteins from intra-vacuolar bacterial pathogens and a putative C-terminal E-block domain (37), suggesting that CstK is indeed a *Coxiella* effector protein (Fig. 1A). Consistently, previous studies by Chen and colleagues have shown that CstK is secreted in a T4SS-dependent manner by the surrogate host *L. pneumophila*, albeit with low efficiency. In order to validate CstK secretion in *C. burnetii*, we engineered plasmids encoding, either CstK or CvpB (a known *C. burnetii* effector protein) (38), fused to Beta-Lactamase (BLAM) and expressed in *Coxiella Tn1832* (a *C. burnetii* transposon mutant that phenocopies wild-type *C. burnetii*) or *dotA*::Tn (a *dotA* transposon mutant, defective for Dot/Icm secretion). By means of a BLAM secretion assay, we could observe that BLAM-CstK was secreted by *Coxiella* at 48 and 72 hours, but not at 24 hours post-infection (Fig. 1B). Secretion of BLAM-CvpB or BLAM-CstK was not detectable in cells infected with the *dotA*::Tn strain, indicating that both CvpB and CstK are *C. burnetii* Dot/Icm substrates (Fig. 1B). Next, we ectopically expressed HA-tagged CstK in non-infected and *C. burnetii* infected U2OS cells, to investigate its intracellular localization. CstK mainly localized at vesicular compartments in non-infected cells, and accumulated at CCVs upon *C. burnetii* infection, suggesting an active role in the biogenesis of this compartment (Figure 1C).

### CstK displays autokinase and protein kinase activities

To determine whether CstK is a functional protein kinase, this protein was overproduced in *E. coli* and purified as a recombinant protein fused to glutathione *S*-transferase (GST) tag. The purified tagged CstK protein (Fig. 2A, upper panel) was then assayed for autokinase activity in presence of the phosphate donor [γ-^33^P]ATP. As shown in Fig. 2A (lower panel), CstK incorporated radioactive phosphate from [γ-^33^P]ATP, generating a radioactive signal corresponding to the expected size of the protein isoform, strongly suggesting that this kinase undergoes autophosphorylation.

**Figure 2.**
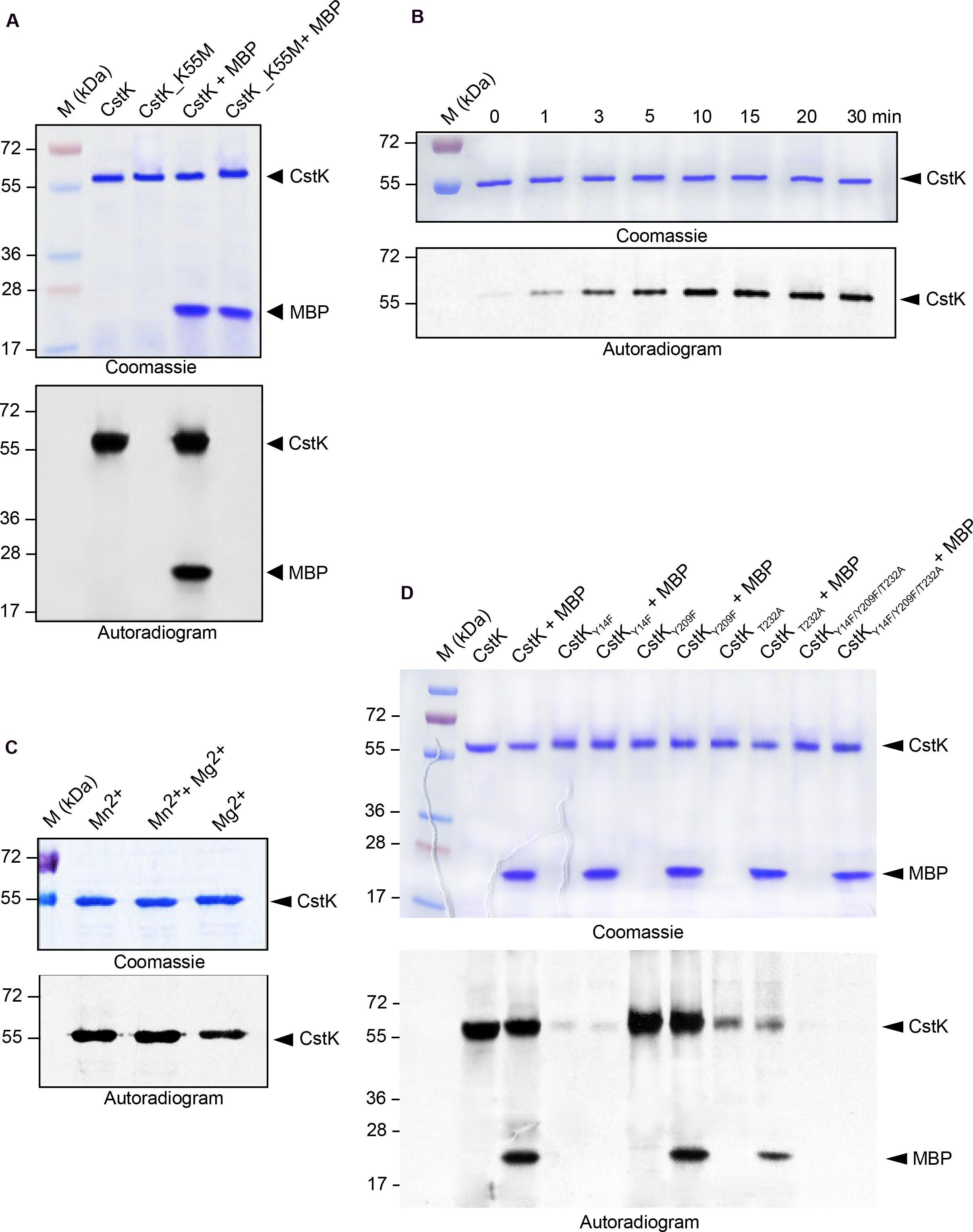
(A) Biochemical characterization of CstK. Recombinant CstK derivatives were overproduced, purified on glutathione-Sepharose 4B matrix and submitted to gel electrophoresis and stained with Coomassie Blue (upper panel). *In vitro* phosphorylation assays were performed with [γ-^33^P]ATP for 30 min and the eukaryotic substrate myelin basic protein (MBP) when required. Proteins were analyzed by SDS-PAGE, and radioactive bands were revealed by autoradiography (lower panel). The upper bands illustrate the autokinase activity of CstK, and the lower bands represent phosphorylated MBP. Standard proteins of known molecular masses were run in parallel. **(B)** A kinetic analysis was performed by incubation of CstK with [γ-^33^P]ATP over different times. Proteins were analyzed by SDS-PAGE, and radioactive bands were revealed by autoradiography. **(C)** Effects of cations on autokinase activity of CstK. Purified CstK protein was subjected to *in vitro* autophosphorylation assays in the presence of [γ-^33^P]ATP and Mg^2+^ or Mn^2+^. Phosphoproteins were separated by SDS-PAGE and then revealed by autoradiography. **(D)** *In vitro* phosphorylation of CstK mutant derivatives. The mutated variants CstK_Y14F (harboring a Tyr to Phe substitution at Y14), CstK_Y209F (harboring a Tyr to Phe substitution at Y209), CstK_T232A (harboring a Thr to Ala substitution at T232), or CstK_Y14F/Y209F/T232A (harboring all three substitutions) were incubated in presence of [γ-^33^P]ATP with or without the eukaryotic substrate myelin basic protein (MBP). Samples were separated by SDS-PAGE, stained with Coomassie Blue (upper panel), and visualized by autoradiography (lower panel). The upper bands illustrate the autokinase activity of CstK, and the lower bands represent phosphorylated MBP. Standard proteins of known molecular masses were run in parallel (kDa lane).

To confirm CstK autophosphorylation and exclude that contaminant kinase activities from *E. coli* extracts might phosphorylate CstK, we mutated the conserved Lys55 residue present in subdomain II into CstK by site-directed mutagenesis. Indeed protein sequence analysis revealed that Lys55 in CstK is similarly positioned as a conserved Lys residue usually involved in the phosphotransfer reaction and also required for the autophosphorylating activity of eukaryotic-like STPKs (34,35). Thus, Lys55 was substituted by a Met residue, and the mutated form of CstK, CstK_K55M, was purified as described above (Fig. 2A, upper panel) and then tested for autophosphorylation in presence of [γ-^33^P]ATP. As expected, no radioactive signal could be detected (Fig. 2A, lower panel), thus establishing that CstK displayed autophosphorylation activity.

A kinetic analysis of CstK phosphorylation was next performed to determine the initial CstK phosphorylation rate (Fig. 2B). Incorporation of γ-phosphate occurred rapidly, reaching about 50% of its maximum rate within 5 min of reaction. This autokinase activity was dependent on bivalent cations such as Mg^2+^ and Mn^2+^ as shown in Fig. 2C, and abolished by addition of 20 mM EDTA (data not shown).

The recombinant CstK protein was further characterized by studying its ability to phosphorylate exogenous proteins and was thus assayed for *in vitro* phosphorylation of the general eukaryotic protein kinase substrate, myelin basic protein (MBP), in the presence of [γ-^33^P]ATP. MBP is a commonly used substrate for both Ser/Thr and Tyr kinases. A radiolabeled signal at the expected 18-kDa molecular mass of MBP was detected, thus demonstrating that CstK phosphorylates protein substrates such as MBP (Fig. 2A). Altogether, these data indicate that *in vitro*, CstK possesses intrinsic autophosphorylation activity and displays kinase functions for exogenous substrates.

### Identification of CstK autophosphorylation sites

To determine the specificity of this kinase, we next identified its autophosphorylation sites. A mass spectrometry approach was used since this technique allows precise characterization of post-translational modifications including phosphorylation (39,40). NanoLC/nanospray/ tandem mass spectrometry (LC-ESI/MS/MS) was applied for the identification of phosphorylated peptides and for localisation of phosphorylation sites in CstK. This approach led to 97% of sequence coverage, while the remaining residues uncovered did not include Ser, Thr or Tyr residues able to be phosphorylated.

As detailed in Table 3, analysis of tryptic digests allowed the characterization of three phosphorylation sites in CstK. Surprisingly, unlike classical Ser/Thr or Tyr kinases, CstK was phosphorylated on two Tyr residues (Y14 and Y209) in addition to one Thr site (T232). Since protein sequence analysis did not reveal a classical activation loop in this kinase, the contribution of T232, Y14 and Y209 to CstK kinase activity was individually assessed. Hence, these residues were mutated either to phenylalanine to replace tyrosine residues or alanine to replace threonine residue, thus generating the single mutants CstkK_Y14F, CstK_Y209F, and CstK_T232A as well as the CstK_Y14F/Y209F/T232A triple mutant. Next, *in vitro* kinase assays with [γ-^33^P]ATP were carried out and revealed that maximum loss in CstK autophosphorylation activity was observed in the CstK_Y14F mutant (Fig. 2D), suggesting that this site is central for CstK activation. In contrast, the CstK_Y209F mutant exhibited a slight hyperphosphorylation, which might indicate that Y209 only plays an accessory role in controlling CstK autophosphorylation (Fig. 2D). Finally, the CstK_T232A mutant showed a reduced CstK phosphorylation and displayed diminished kinase activity towards the exogenous substrate MBP (Fig. 2D). Note that mutating all three autophosphorylation sites fully abrogated CstK kinase activity (Fig. 2D). These results indicate that Y14 and T232 are the major phosphorylation sites in CstK and strongly suggest that CstK might be a dual specificity (Thr/Tyr) kinase.

### CstK regulates vacuole development and C. burnetii replication within infected cells

As a first step towards the understanding of CstK functions in the course of infection, and to appreciate the extent to which this kinase is required for growth and viability of *C. burnetii*, we attempted to inactivate the corresponding chromosomal gene. Unfortunately, after several attempts we were unable to generate a null mutant suggesting that *cstK* might be essential. However, we had previously isolated a *C. burnetii* mutant (*Tn2496*) carrying a transposon insertion at position 156783, 32 bp upstream of the starting codon of *cstK* (41) (Fig. 1A). To determine the effect of this transposon insertion for *cstK* gene expression, we assessed the expression level of *cstK* mRNA from the *C. burnetii* wild type and *Tn2496* strains. Surprisingly, *cstK* expression was significantly upregulated in the mutant strain, suggesting that the transposon insertion may have released a transcriptional negative regulation (Fig. 3A). This suggested that a putative transcriptional regulator might bind the *cstK* promoter and control its activity during host invasion.

**Figure 3.**
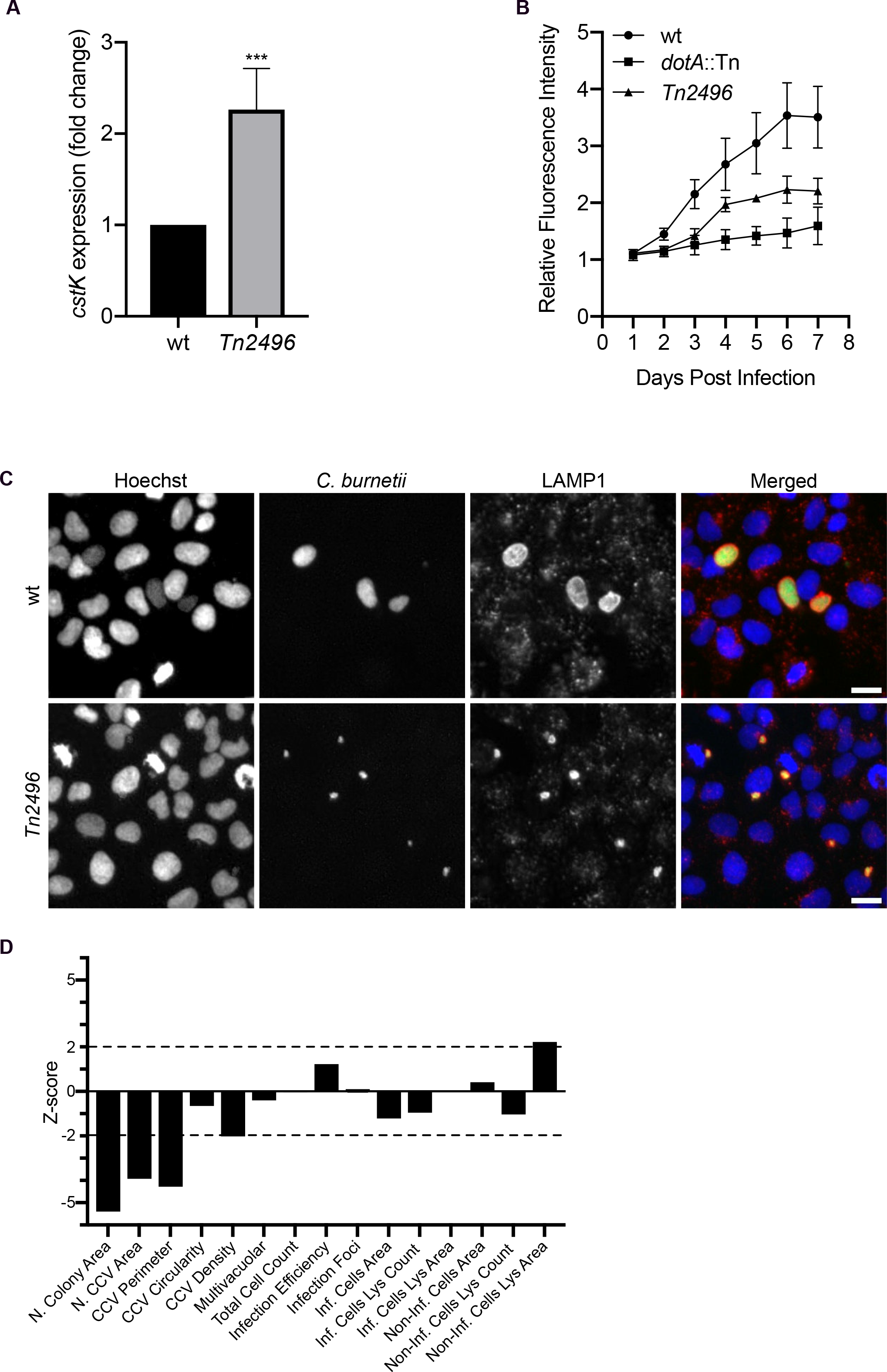
CstK regulates vacuole development and *C. burnetii* replication within infected cells. **(A)** expression of *cstk* was measured by qRT-PCR from a culture of *C. burnetii* Tn1832 (wt) strain and *C. burnetii* mutant *Tn2496*. Values are means ± SEM of 4 independent experiments. ***, *p* < 0.001, unpaired t-test. **(B)** Vero cells were challenged with either *C. burnetii* wild-type (wt), the *dotA*::Tn mutant or with the mutant *Tn2496*. Intracellular replication was monitored over 7 days of infection using a microplate reader exploiting the GFP expressed by *C. burnetii* strains. Values are mean ± SD from 3 independent experiments. **(C)** U2OS cells challenged with either *C. burnetii Tn1832* (wt) or *Tn2496* were fixed and labelled with an anti-LAMP1 antibody and Hoechst. **(D)** An average of 50000 cells were automatically imaged and analysed for each condition and the phenotypic profile of the *Tn2496* mutant was compared to that of *Tn1832* and expressed as z-scores over 15 independent features.

We next examined the phenotypic effects of CstK overexpression in *C. burnetii*. This mutant was tested for its capacity of invading and replicating within host cells. *Tn2496* maintained the capacity of invading host cells (Fig. 3B), however, intracellular growth was significantly reduced over 7 days of infection with an intermediate phenotype between WT and the non-secreting *dotA*::Tn mutant (Fig. 3B&3C). Accordingly, this was reflected in a strong reduction in the size of intracellular *C. burnetii* colonies and CCVs (Fig. 3D). Therefore, we could conclude that CstK might participate in the formation of the *C. burnetii* replicative vacuole, and that its expression must be finely tuned for optimal intracellular replication.

### CstK specifically interacts with host cell proteins

Since CstK is a secreted protein, we assume that this kinase would interfere with host cell signal transduction pathways to subvert host cell defenses to the benefit of the bacteria. To identify host cell proteins that could interact with CstK, we made use of the model amoeba *Dictyostelium discoideum*. *D. discoideum* is a eukaryotic professional phagocyte amenable to genetic and biochemical studies. Lysate from cells overexpressing CstK tagged with a C-terminal FLAG epitope (CstK-FLAG) was incubated with beads coupled to an anti-FLAG antibody. Beads were extensively washed and bound proteins were separated by SDS-PAGE before mass-spectrometry analysis. The putative interactants of CstK identified by this approach are mainly involved in the endocytic pathway (data not shown).

### CstK interacts with TBC1D5 in vitro and in cells

Among several identified proteins, we focused on a RabGAP/TBC domain-containing protein, DDB_G0280253 (UniProtKB - Q54VM3). This 136.4 kDa protein is homologous to mammalian TBC1D5 (http://dictybase.org), a GTPase-activating protein (GAP) for Rab7a (42), thus an interesting putative target regarding the role of Rab7a in *C. burnetii* vacuole formation (43). Therefore, to verify the interaction between CstK and TBC1D5, we assayed the interaction *in vivo* of FLAG-CstK with human TBC1D5 (Hs-TBC1D5) by using a GFP Trap assay (Fig. 4A). HEK cells expressing FLAG-CstK, Hs_TBC1D5-GFP or both proteins were used for immunoprecipitation using GFP-trap beads and CstK was only detected as co-immunoprecipitated in presence of Hs_TBC1D5-GFP thus confirming that Hs-TBC1D5 is a bona fide CstK interactant *in vivo*, though we cannot formally exclude that other host cell proteins contribute to this interaction. This interaction was also observed in confocal microscopy studies. Ectopic expression of HA-CstK and GFP-TBC1D5 in non-infected U2OS cells, resulted in the recruitment of TBC1D5 at CstK-positive compartments, as opposed to cells overexpressing GFP-TBC1D5 alone, where the TBC1D5 was found diffused in the cytoplasm (Fig. 4B).

**Figure 4.**
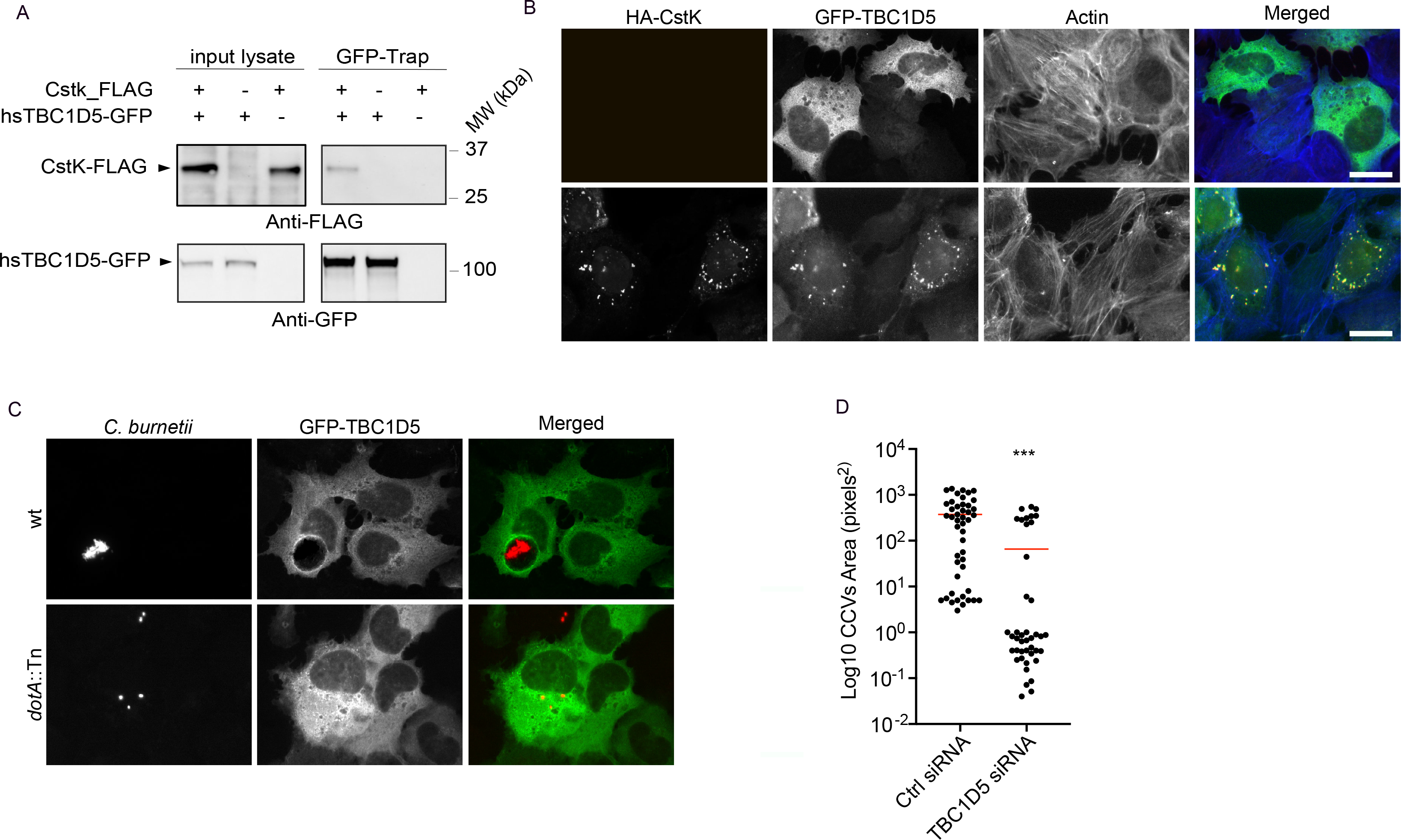
CstK interacts with TBC1D5. **(A)** The interaction between CstK and the human homologue of TBC1D5 has been confirmed after cotransfection of HEK cells to express either a FLAG-tagged CstK, a GFP-tagged Hs_TBC1D5 or both proteins. Cells have been lysed and Hs_TBC1D5-GFP has been trapped on anti GFP beads (GFP Trap-Chromotech). Beads were washed, eluted by boiling, and bound proteins were revealed by Western blot analysis. Anti-GFP antibody (lower panel) confirms the immunoprecipitation of Hs_TBC1D5-GFP. The co-immunoprecipitation of CstK_FLAG is confirmed in presence of Hs_TBC1D5 (upper panel). **(B)** U2OS cells expressing GFP-tagged Hs_ TBC1D5 (green) alone (upper panel), or in combination with HA-tagged CstK (red) were fixed and labelled using Alexa647-labelled phalloidin (blue). Scale bars 10 μm. **(C)** Hs_ TBC1D5 localisation during infection was monitored in U2OS cells expressing GFP-tagged Hs_ TBC1D5 (green) challenged either with *C. burnetii Tn1832* (wt) or *dotA*::Tn (Dot/Icm-defective; red). Scale bars 10 μm. **(D)** The role of TBC1D5 in *C. burnetii* infections was investigated using siRNA to deplete TBC1D5 in U2OS prior to challenge with *C. burnetii Tn1832* (wt). The size of CCVs was automatically calculated over an average of 150000 cells per condition. Red bars indicate medians. ***, *p* < 0.001, one-way ANOVA, Dunnett’s multiple comparison test.

### TBC1D5 is recruited at CCVs and regulates their biogenesis

Given its recently reported role in the development of *L. pneumophila*-containing vacuoles, we investigated the localization of TBC1D5 in U2OS cells infected either with *C. burnetii* control transposon mutant *Tn1832* or the Dot/Icm-defective mutant *dotA*::Tn. TBC1D5 seems to accumulate at CCVs in a Dot/Icm-dependent manner as the eukaryotic protein failed to accumulate around intracellular *C. burnetii dotA*::Tn (Fig. 4C). Next, we used siRNA to deplete cells of TBC1D5 prior to *C. burnetii* infection, to investigate a possible role in CCVs development and intracellular bacterial replication. Indeed, vacuole development was significantly reduced in cells exposed to TBC1D5-targeted siRNAs as opposed to cells treated with non-targeting siRNA oligonucleotides (Fig. 4D).

### TBC1D5 is not phosphorylated in vitro by recombinant CstK

Next, we assessed whether CstK might phosphorylate the recombinant Hs-TBC1D5. Despite the *in silico* prediction of several Ser/Thr and Tyr phosphorylatable residues in Hs-TBC1D5, we failed to detect Hs-TBC1D5 phosphorylation using several *in vitro* kinases assays (data not shown). In addition, Hs-TBC1D5 phosphorylation status was also investigated upon transfection with CstK or its inactive derivative (K55M) followed by Hs-TBC1D5 immunoprecipitation. No phosphorylation could be detected in our conditions, though a sensitivity issue regarding the detection cannot be ruled out.

## DISCUSSION

Our results provide the first biochemical analysis of the secreted kinase CstK and its involvement in the process of infection and CCVs development. Our study also delineates important differences with the classical Ser/Thr kinases. In particular, we provided evidence that CstK is a dual kinase able to autophosphorylate on Thr and Tyr residues. Our overexpressing mutant suggested the presence of a negative transcriptional regulation of *cstK* expression and its phenotype highlighted the importance of regulating *cstK* expression during *C. burnetii* infections. Moreover, the identification of candidate host interactors of CstK further corroborated an important role of the bacterial kinase during infection. We were not able to detect TBC1D5 phosphorylation by CstK in our conditions. However, lack of phosphorylation of host interactors of bacterial STPKs is not uncommon (21) as interaction between STPKs and host proteins might as well perturb protein interaction networks at play in host cells. The biochemical mechanisms of these pathogen-directed targeted perturbations of host cell-signaling networks are being actively investigated. Regardless, siRNA depletion of TBC1D5 in *C. burnetii*-infected cells points at a role of the eukaryotic protein in CCVs development. In mammals, TBC1D5 was suggested to function as a molecular switch between endosomal and autophagy pathways. Indeed TBC1D5 associates the retromer VPS29 subunit involved in endosomal trafficking, and upon autophagy induction, the autophagy ubiquitin-like protein LC3 can displace VPS29, thus orienting TBC1D5 functions towards autophagy instead of endosomal functions (44). It is thus tempting to propose that CstK might interfere with this tight regulation between TBC1D5, LC3 and VPS29, and redirect TBC1D5 functions to support efficient *C. burnetii* intracellular replication. Further work needs to be carried out to understand how CstK recognize these host substrates and how they participate in the establishment of *C. burnetii* parasitophorous vacuoles. Another perspective of this work is the opening of a new field of investigation for future drug development to fight this pathogen. Because CstK seems to be essential, specific inhibitors capable of preventing *C. burnetii* growth would be extremely useful for the development of new therapies.

## EXPERIMENTAL PROCEDURES

### Bacterial strains and growth conditions

Bacterial strains and plasmids are described in Table 1. Strains used for cloning and expression of recombinant proteins were *Escherichia coli* TOP10 (Invitrogen) and *E. coli* BL21(DE3)Star (Stratagene), respectively. *E. coli* cells were grown and maintained at 25 °C in LB medium supplemented with 100 µg/ml ampicillin when required. *Coxiella burnetii* RSA439 NMII and transposon mutants *Tn1832*, *Tn2496*, and *dotA::Tn* were grown in ACCM-2 (45) supplemented with kanamycin (340 mg/ml) or chloramphenicol (3 mg/ml) as appropriate in a humidified atmosphere of 5% CO2 and 2.5% O2 at 37°C.

**Table 1.** Strains and plasmids used in this study

| Strains or Plasmids | Genotype or Description <sup>a</sup> | Source or Reference |
| --- | --- | --- |
| ***Escherichia coli* strains** |  |  |
| *E. coli* 10G | *E. coli* derivative ultra-competent cells used to general cloning; F- mcrA D(mrr-hsdRMS-mcrBC) f80dlacZΔM15 D lacX74 endA1 recA1araD139 D (ara, leu)7697 galU galK rpsL nupG - tonA | Lucigen |
| *E. coli* BL21(DE3)Star | F2 ompT hsdSB(rB2 mB2) gal dcm (DE3); used to express recombinant proteins in *E. coli* | Stratagene |
| ***Coxiella burnetii* strains** |  |  |
| GFP-*Coxiella burnetii* RSA439 NMII | *Coxiella burnetii* RSA439 NMII carrying a Tn7-CAT-GFP cassette in the intergenic region between CBU_1787 and CBU_1788 | (50) |
| *Coxiella burnetii* Tn2496 | *Coxiella burnetii* RSA439 NMII carrying an Himar1-CAT-GFP cassette 32 bp upstream of CBU_0175 | This study |
| *Coxiella burnetii* Tn1832 | *Coxiella burnetii* RSA439 NMII carrying an Himar1-CAT-GFP cassette in in the intergenic region between CBU_1847b and CBU_1849 | (49) |
| *Coxiella burnetii* Tn292 | *Coxiella burnetii* RSA439 NMII carrying an Himar1-CAT-GFP cassette in CBU_1648 (*dotA*) | (49) |
| **Plasmids** |  |  |
| pGEX(M) | pGEX with a 321-bp *EcoRI/BamHI* fragment from pET19b introducing a *HindIII* site in the pGEX polylinker | (51) |
| pGEX(M)_cstK | pGEX(M) derivative used to express GST-tagged fusion of CstK (Amp<sup>R</sup>) | This study |
| pGEX(M)_cstK_K55M | pGEX(M) derivative used to express GST-tagged fusion of CstK_K55M (Amp<sup>R</sup>) | This study |
| pGEX(M)_cstK_Y14F | pGEX(M) derivative used to express GST-tagged fusion of CstK_Y14F (Amp<sup>R</sup>) | This study |
| pGEX(M)_cstK_Y209F | pGEX(M) derivative used to express GST-tagged fusion of CstK_Y209F (Amp<sup>R</sup>) | This study |
| pGEX(M)_*cstK*_T232A | pGEX(M) derivative used to express GST-tagged fusion of CstK_T232A (Amp<sup>R</sup>) | This study |
| pGEX(M)_*cstK*_Y14F/Y209F/T232A | pGEX(M) derivative used to express GST-tagged fusion of CstK_Y14F/Y209F/T232A (Amp<sup>R</sup>) | This study |
| pcDNA3.1(-)_Flag_*cstK* | pcDNA3.1(-) derivative used to express N-terminal Flag-tagged fusion of CstK for mammalian expression | This study |
| peGFPN1_Hs-TBC1D5-GFP | peGFP-N1 derivative used to express C-terminal GFP-tagged fusion of Hs_TBC1D5 for mammalian expression | This study |
| pRK5_HA_*cstK*opt | pRK5_HA derivative used to express HA-tagged fusion of CstK codon-optimized for mammalian expression | This study |
| pXDC61K-Blam | Vector with IPTG-inducible expression of Beta-Lactamase (Blam) | (52) |
| pXDC61K-Blam-*cvpB* | pXDC61K-Blam derivative used to express Blam-tagged fusion of CvpB | This study |
| pXDC61K-Blam-*cstK* | pXDC61K-Blam derivative used to express Blam-tagged fusion of CstK | This study |
| ***Dictyostelium discoideum* plasmids** |  |  |
| pDXA-3C_*cstK*-FLAG | vector for overexpression of CstK with a C-terminal Flag tag in *D. discoideum* | This study |
| pDXA-3C_*cstK*-myc | vector for expression of myc-DdTBC1D5 in *D. discoideum* | This study |

<sup>a</sup> Amp<sup>R</sup>, ampicillin resistant; Cm<sup>R</sup>, chloramphenicol resistant; Km<sup>R</sup>, kanamycin resistant; Spec<sup>R</sup>, spectinomycin resistant; Tet<sup>R</sup>, tetracycline resistant

### Cloning, expression and purification of CstK derivatives

The *cstK (CBU_0175)* gene was amplified by PCR using *Coxiella burnetii* RSA439 NMII chromosomal DNA as a template with the primers listed in Table 2 containing a *Bam*HI and *Hind*III restriction site, respectively. The corresponding amplified product was digested with *Bam*HI and *Hind*III, and ligated into the bacterial pGEX(M) plasmid, that includes a N-terminal GST-tag, thus generating pGEX(M)_*cstK.* pGEX(M)_*cstK* derivatives harboring different mutations were generated by using the QuikChange Site-Directed Mutagenesis Kit (Stratagene, La Jolla, CA), and resulted in the construction of plasmids detailed in Table 2. All constructs were verified by DNA sequencing. Purifications of the GST-tagged recombinants were performed as described by the manufacturer (GE Healthcare). For expression in mammalian cells, the *cstK* gene was amplified by PCR using pGEX(M)_*cstK* as a template with primers listed in Table 2 containing a *Bam*HI and *Hind*III restriction site, respectively. The corresponding amplified product was digested with *Bam*HI and *Hind*III, and ligated into the eukaryotic pcDNA3.1(-) plasmid, thus generating pcDNA3.1(-)_Flag_*cstK. cstK* coding sequence was also optimized for mammalian cell expression (GenScript), amplified by PCR and cloned into pDXA-3C (46) containing a FLAG-tag for C-terminal fusion. After sequencing, the plasmid was linearized by the restriction enzyme *Sca*I and transfected in *D. discoideum* as described (47). Clone selection was made with 10 mg/ml of G418, and protein expression was assayed by Western blot analysis of *D. discoideum* crude extract with an anti-FLAG rabbit polyclonal antibody (GenScript, USA). For ectopic expression assays, *cstK* with optimized codons was cloned in pRK5-HA using the primer pair CstKopt-BamHI-Fw/CstKopt-EcoRI-Rv.

**Table 2.**
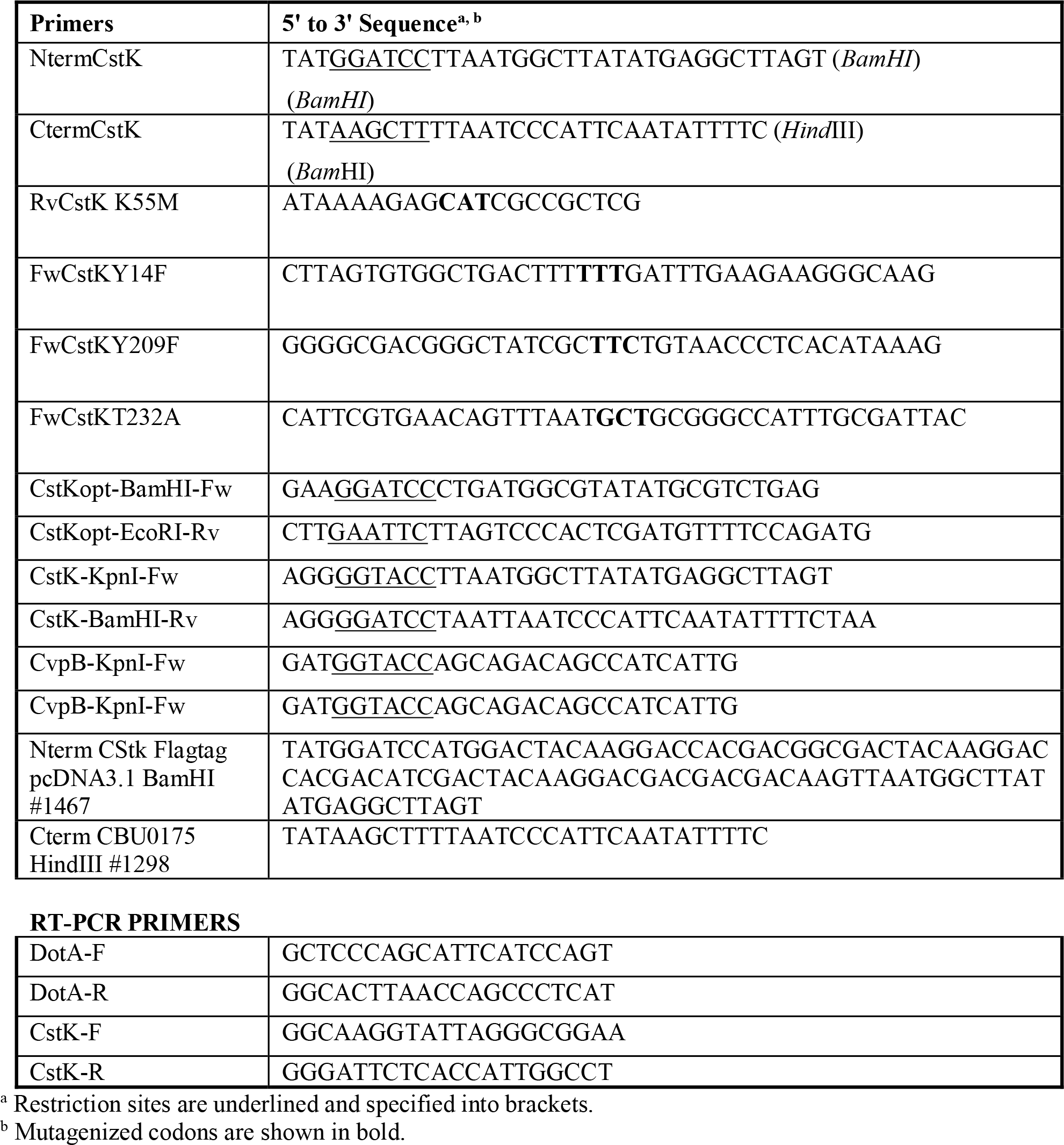
Primers used in this study

**Table 3.**
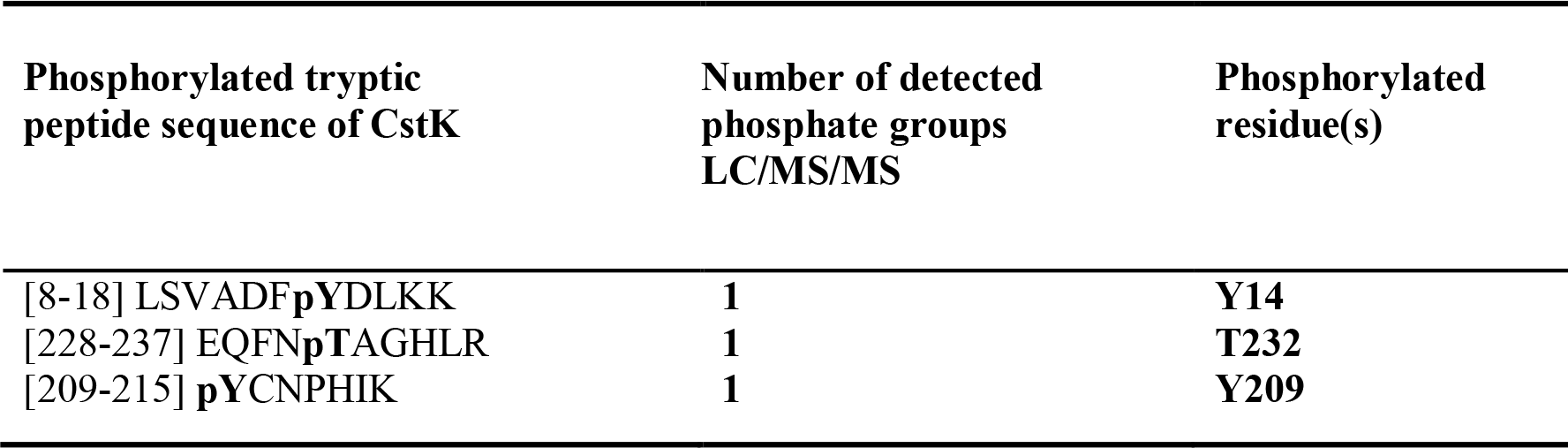
Phosphoacceptors identified after phosphorylation of *C. burnetii* CstK. Sequences of the phosphorylated peptides identified in CstK as determined by mass spectrometry following tryptic digestion are indicated, and phosphorylated residues (pT or pY) are shown in bold.

| Phosphorylated tryptic peptide sequence of CstK | Number of detected phosphate groups LC/MS/MS | Phosphorylated residue(s) |
| --- | --- | --- |
| [8-18] LSVADF**p**YDLKK | **1** | **Y14** |
| [228-237] EQFN**p**TAGHLR | **1** | **T232** |
| [209-215] **p**YCNPHIK | **1** | **Y209** |

### Cloning, expression and purification of TBC1D5 derivatives

The *D. discoideum* GFP-tagged TBC1D5 was previously generated (48). Cells were grown at 22°C in HL5 medium as previously described (47). Human TBC1D5 coding sequence was obtained from the hORFeome v8.1 (ORF 2659, Q92609, fully-sequenced cloned human ORFs in Gateway Entry clones ready for transfer to Gateway-compatible expression vectors). HsTBC1D5 coding sequence has been recombined into pEGFP-N1 RfC Destination vector by GateWay reaction (MGC Paltform Montpellier) thus generating pEGFP-N1_HsTBC1D5 coding for HsTBC1D5 with a C-terminal GFP Tag.

### RNA extraction and quantitative RT-PCR (qRT-PCR)

Fifty ml of *C. burnetii* culture was harvested, resuspended in 600 µL of RNA protect reagent (Qiagen) and incubated 5 min at room temperature. Bacteria were centrifuged and resuspended in 200 µl TE (10 mM Tris-HCl, 1 mM EDTA, pH 8) containing 1 mg/ml lysozyme. Bacterial suspension was incubated at room temperature for 5 min and bacteria were disrupted by vigorous vortexing for 10 sec every 2 min. 700 µl of RLT buffer from RNA easy kit (Qiagen), were added to the bacterial suspension and disruption was completed by vortexing vigorously. RNA was purified with the RNA easy kit according to manufacturer’s instructions. DNA was further removed using DNAse I (Invitrogen). cDNA was produced using Superscript III reverse transcriptase (Invitrogen). Controls without reverse transcriptase were done on each RNA sample to rule out possible DNA contamination. Quantitative real-time PCR was performed using LightCycler 480 SYBR Green I Master (Roche) and a 480 light cycler instrument (Roche). PCR conditions were as follows: 3 min denaturation at 98°C, 45 cycles of 98°C for 5 sec, 60°C for 10 sec and 72°C for 10 sec. The *dotA* gene was used as internal control. The sequences of primers used for qRT-PCR are listed in Table 2. The expression level of *cstK* in the wild type strain was set at 100 and the expression levels of *cstK* in the *Tn2496* mutant were normalized to the wild type levels.

### In vitro kinase assays

*In vitro* phosphorylation was performed with 4 μg of wild-type CstK or CstK derivatives in 20 µl of buffer P (25 mM Tris-HCl, pH 7.0; 1 mM DTT; 5 mM MnCl_2_; 1 mM EDTA; 50μM ATP) with 200 µCi ml^−1^ (65 nM) [γ-^33^P]ATP (PerkinElmer, 3000 Ci mmol^−1^) for 30 min at 37 °C. For substrate phosphorylation, 4 µg of MBP (Myelin Basic Protein) (Sigma) and 4 µg of CstK were used. Each reaction mixture was stopped by addition of an equal volume of Laemmli buffer and the mixture was heated at 100 °C for 5 min. After electrophoresis, gels were soaked in 16% TCA for 10 min at 90 °C, and dried. Radioactive proteins were visualized by autoradiography using direct exposure to films.

### Mass spectrometry analysis

For mass spectrometry analysis, CstK was phosphorylated as described above, excepted that [γ-^33^P]ATP was replaced with 5 mM cold ATP. Subsequent mass spectrometry analyses were performed as previously reported (31–34). Spectra were analyzed with the paragon algorithm from the ProteinPilot® 2.0 database-searching software (Applied Biosystems) using the phosphorylation emphasis criterion against a homemade database that included the sequence of CstK.

### C. burnetii infections

U2OS epithelial cells were challenged either with *Coxiella burnetii* RSA439 NMII, the transposon mutants *Tn1832*, *dotA*::Tn or *Tn2496* as previously described (38,49). For gene silencing, U2OS cells were seeded at 2,000 cells per well in black, clear-bottomed, 96-well plates in triplicate and transfected with siRNA oligonucleotides 24 h later by using the RNAiMAX transfection reagent (Thermo Fisher Scientific) according to the manufacturer’s recommendations. At 24 h post transfection, cells were challenged with *C. burnetii* (MOI of 100) and further incubated for 5 d. Cells are then fixed and processed for immunofluorescence. Samples were imaged with an ArrayScan VTI Live epifluorescence automated microscope (Cellomics) equipped with an ORCA-ER CCD camera (Hamamatsu). 25 fields per well were acquired for image analysis. ImageJ, ICY, and CellProfiler software were used for image analysis and quantifications. Phenotypic profiles (expressed as z-scores) were calculated following median based normalization of 96-well plates. Plates effects were corrected by the median value across wells that are annotated as control.

### Beta-lactamase translocation assay

Effector proteins translocation assays were performed as previously described (38). Briefly, *Coxiella burnetii Tn1832* (wt) and *dotA*::Tn were transformed either with pXDC-Blam (negative control), pXDC-Blam-CvpB (positive control) or pXDC-Blam-CstK. Each strain was used to infect U2OS epithelial cells. After 24, 48 or 72 hours of infection, cells were loaded with the fluorescent substrate CCF4/AM (LiveBLAzer-FRET B/G loading kit; Invitrogen) in HBSS (20 mM HEPES pH 7.3) containing 15 mM probenecid (Sigma). Cells were incubated in the dark for 1 h at room temperature and imaged using an EVOS inverted fluorescence microscope. Images were acquired using DAPI and GFP filter cubes. The image analysis software CellProfiler was used to segment and identify all cells in the sample (GFP), positive cells (DAPI) and to calculate the intensity of fluorescence in each channel. The percentage of positive cells versus the total number of infected cells was then calculated and used to evaluate effector translocation.

### Immunoprecipitation from D. discoideum lysates

For immunoprecipitation assays, 2×10^7^ cells were lysed in lysis buffer (50 mM Tris-HCl pH 7.5, 300 mM NaCl, 0.5% NP40, protease inhibitors (Roche)) and cleared by centrifugation for 15 min at 14 000 rpm in a microfuge. Lysate supernatants were incubated overnight at 4°C with anti-flag monoclonal antibody coated on agarose beads (Genscript). The beads were then washed five times in wash buffer (50mM Tris-HCl pH 7.5, 300 mM NaCl, 0,1% NP40) and once in PBS. Bound proteins were migrated on SDS-PAGE and analysed by LC-MS/MS.

### GST pull-down

GST fusion proteins were produced as described above and bound to Glutathione Sepharose 4B beads according to manufacturer’s instructions (GE Heathcare). To prepare cell lysates, *D. discoideum* cells (2×10^7^) were incubated 15 min in lysis buffer (20 mM HEPES buffer, pH 7.0; 100 mM NaCl; 5 mM MgCl_2_; 1% Triton X100; protease inhibitors (Roche)) and centrifuged for 15 min at 14 000 rpm in a refrigerated microfuge. GST beads were then incubated with cell lysates (800 µg) overnight at 4°C on a wheel. After three washes in lysis buffer, beads were heated at 95°C for 10 min. Bound proteins were separated by SDS-PAGE and transferred to nitrocellulose before incubation with an anti-myc antibody (9E10; Sigma). Blots were revealed with the Odyssey Western Detection System.

### Cell culture, heterologous expression and GFP-Trap immunoprecipitation

HEK cells were grown in a DMEM (Gibco) containing 10% (vol/vol) FBS, 1% glutamax (Gibco, 200 mM stock), 0,5% Penicillin/Streptomycin (Gibco, 10000 U/mL stock) and maintained under standard conditions at 37 °C in a humidified atmosphere containing 5% CO2. Cells were transiently transfected using the jetPEI Transfection Reagent (Polyplus-Transfection Inc.) to express either Hs-TBC1D5_GFP, CstK_FLAG or both proteins. Cells were used 48 h after transfection for immunoprecipitation assay. Transfected cells were washed two times in cold PBS and lysed in lysis buffer (20 mM Hepes, 100 mM NaCl, 5 mM MgCl_2_, 0,2 % Triton X100, 10 % Glycerol, protease inhibitors (Roche)). Cleared lysate (700 µL, approximately 0.8 mg total proteins) were incubated with GFP-Trap (Chromotek) for 30 min at room temperature under gentle rotation. The beads were washed three times in lysis buffer, boiled in 2X Laemmli sample buffer and loaded on ExpressPlus™ PAGE Gels (GenScript). The eluted proteins were visualized by Western Blotting with the following antibodies: anti-FLAG M2 from Sigma Aldrich, anti-GFP from Torrey Pines, donkey anti-mouse or anti-rabbit from Jackson ImmunoResearch.

## ACKNOWLEDGMENTS

The wish to thank the Montpellier RIO imaging facility at the University of Montpellier 2. This work was supported by grants from the ATIP/AVENIR Program for V.M. and M. B., the Region Languedoc-Roussillon for S. B., a Marie Curie CIG to E.M. (grant n. 293731), the Agence Nationale de la Recherche (ANR) Grant ANR-14-CE14-0012-01 (project AttaQ) to M.B.

